# Adversary of DNA integrity: a long non-coding RNA stimulates driver oncogenic chromosomal rearrangement in human thyroid cells

**DOI:** 10.1101/2022.03.25.485761

**Authors:** Denis Eriksonovich Demin, Matvey Mikhailovich Murashko, Aksinya Nicolaevna Uvarova, Ekaterina Mikhailovna Stasevich, Elena Yurievna Shyrokova, Gennady Efimovich Gorlachev, Kirill Viktorovich Korneev, Alina Sergeevna Ustiugova, Elena Andreevna Tkachenko, Valentina Vitalevna Kostenko, Karina Aleksandrovna Tatosyan, Saveliy Andreevich Sheetikov, Pavel Vladimirovich Spirin, Dmitriy Vladimirovich Kuprash, Anton Markovich Schwartz

**Author notes:** These authors contributed equally to the manuscript (Denis Eriksonovich Demin, Matvey Mikhailovich Murashko).

## Abstract

The flurry of publications devoted to the functions of long non-coding RNAs (lncRNAs) published in the last decade leaves no doubt about the exceptional importance of lncRNAs in various areas including tumor biology. Contribution of lncRNAs to the early stages of oncogenesis remains poorly understood. In this study we explored a new role for lncRNAs: stimulation of driver oncogenic mutations that result from specific chromosomal rearrangements. We demonstrated that lncRNA CASTL1 (ENSG00000269945) stimulates the formation of the CCDC6-RET inversion (RET/PTC1) in human thyroid cells subjected to radiation or chemical DNA damage. Facilitation of chromosomal rearrangement requires lncRNA to contain regions complementary to the introns of both CCDC6 and RET genes as deletion of these regions deprives CASTL1 of the ability to stimulate the gene fusion. We found that CASTL1 expression is elevated in tumors with CCDC6-RET fusion which is the most frequent rearrangement in papillary thyroid carcinoma. Our results open a new venue for the studies of early oncogenesis in various tumor types, especially those associated with physical or chemical DNA damage.

**Graphical abstract:** 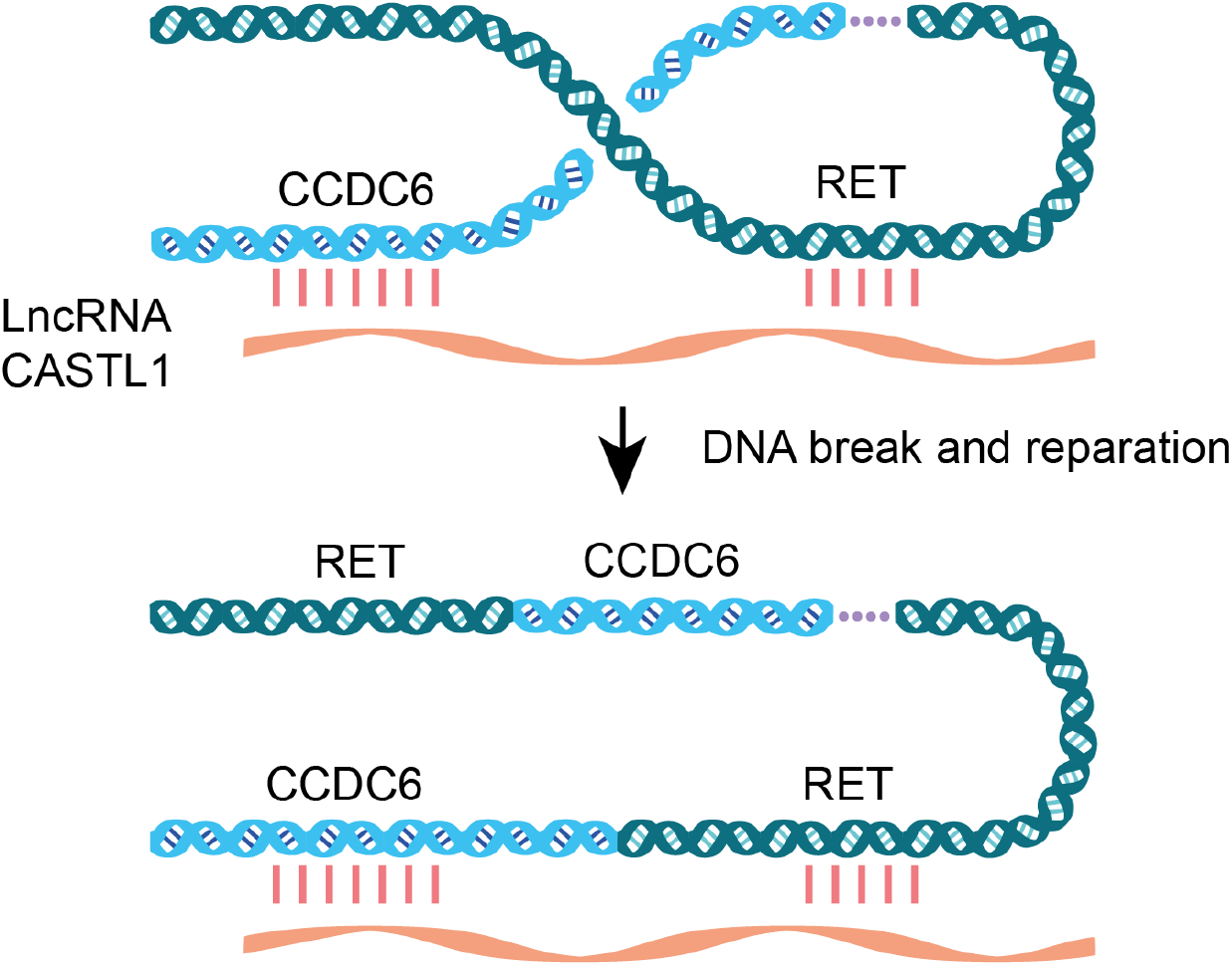

## Introduction

Chromosomal rearrangements are one of the major factors responsible for the formation of driver mutations in various types of tumors ^1,2^. Interestingly, many recurrent chromosomal rearrangements are specific for tumors of different tissues and organs ^3^. The reason for this phenomenon remains poorly understood. A pioneer study has recently demonstrated that expression of CEP131 gene mRNA in a human prostate cell line stimulates the formation of the TMPRSS2-ERG chromosomal aberration characteristic of prostate tumors ^4^. This mRNA contains sequences complementary to the fused genomic regions. It is important to note that the complementary sites of RNA and introns were located near the microhomology region of the fused genes. Of note, in order to stimulate the aberration, RNA should be complementary to the coding strand of the fused genes. Although RNA complementary to the template strand can also stimulate rearrangement when transcription is suppressed ^4^. Gupta et al suggested that as RNA polymerase II moves along the DNA template strand, it throws off the “sense” chimeric RNA during transcription, whereas the “antisense” RNA binds to the DNA coding strand and is not in the path of the moving polymerase. The authors showed that an RNA-DNA interaction site with a length of only 20 nt is sufficient to stimulate gene fusion.

Here, we hypothesized that long non-coding RNAs (lncRNA) may play a similar role. Moreover, the contribution of these RNAs to the stimulation of various chromosomal rearrangements may be higher than that of mRNAs, for a number of reasons. Firstly, unlike mRNAs that are transported from the nucleus into the cytoplasm after synthesis, lncRNAs preferentially localize in the nucleus ^5^. Secondly, lncRNAs tend to be more tissue-specific than protein-coding RNAs ^5,6^. Finally, the sequences of most lncRNAs contain repetitive elements (REs) such as LINE, SINE, and LTR, which are also frequently found in introns ^7,8^. Thus, transcripts capable of complementary interactions with sequences of two different genomic loci are more likely to be found among lncRNAs.

Possible involvement of lncRNA in the stimulation of chromosomal rearrangements between the regions containing REs is consistent with the increased frequency of aberrations between such regions ^9,10^. This phenomenon can be explained by the ability of some REs to facilitate DNA breaks ^11^, as well as by the false recognition of repetitive sites as regions of homology during reparation. However, a recent study shows that chromosomal rearrangements occur less frequently between regions containing two repeats of the AluJ family than between regions containing repeats of two different families, AluY and AluS ^10^. This may imply an alternative mechanism other than the degree of homology between REs that determines the mutation frequency. One such mechanism may be implemented via lncRNAs.

The involvement of lncRNAs in the formation of certain chromosomal rearrangements in eukaryotes has been extensively studied in the case of Ciliates where lncRNAs are involved in the reorganization of the genome during macronucleus formation. These RNAs determine which parts of the genome are excised and which should be joined and in what order ^12^. In human cells, a number of studies have previously reported that lncRNAs are involved in the regulation of reparation and nascent RNAs can control the fidelity of this process via complementary interactions with the joining DNA fragments ^13,14^. This evidence, albeit sparse at the moment, indicates that lncRNAs are indeed involved in chromosomal rearrangements in humans.

For this study, we selected the inversion inv(10)(q11;q21), leading to CCDC6-RET oncogene formation by joining introns 1 and 11 of CCDC6 and RET genes, respectively ^15^. This fusion is the most frequent aberration in papillary thyroid carcinoma ^16–18^. CCDC6-RET belongs to the class of RET/PTC rearrangements, which are found in the majority of patients with thyroid cancer younger than 10 years ^19^. This rearrangement also occurs in patients with lung adenocarcinoma ^20^, colorectal cancer and acute lymphoblastic leukemia ^21,22^. Tumors with the inv(10)(q11;q21) may be resistant to radioiodine treatment (thyroid tumor) and to EGFR inhibitor therapy (lung cancer) ^23,24^. The frequency of this inversion is significantly increased in patients who have been exposed to radiation or received chemotherapy drugs ^16,25^. Also, tumors with the CCDC6-RET oncogene are frequently found in patients with autoimmune disease of the gland ^26^.

In this work, we demonstrate that lncRNA ENSG00000269945.1 (hg38 chr14:71,219,100-71,222,724) (henceforth CASTL1 for <u>c</u>hromosomal <u>a</u>berrations <u>st</u>imulating lncRNA 1), which contains regions complementary to sequences of 1st and 11th introns of CCDC6 and RET genes respectively, stimulates the formation of the chimeric CCDC6-RET oncogene. The discovery of a lncRNA directly involved in the formation of an oncogenic chromosomal rearrangement, particularly upon physical or chemical exposures that cause DNA damage may contribute to the development of novel approaches to cancer therapy.

## Materials and methods

### Cell culture

Nthy-ori 3-1 cell line derived from follicular thyroid epithelium (Sigma - Aldrich, 90011609) was cultured in Cell Culture Treated Flasks at 37°C in the presence of 5% CO2 in RPMI-1640 medium containing 2mM glutamine (Paneco, Russia) supplemented with 10% fetal calf serum (Corning, USA), 10mM HEPES (GIBCO, USA), 1 mM sodium pyruvate (GIBCO, USA), 100 U/ml penicillin and 100 mg/ml streptomycin (Paneco, Russia), and 1x MEM Non-Essential Amino Acids (GIBCO, USA).

### Search for candidate lncRNAs

A search for lncRNAs potentially capable of inducing CCDC6-RET chromosomal rearrangement was performed among non-coding transcripts from the GENCODE database Release 31 ^27^. lncRNAs with regions of homology to intron 1 of the CCDC6 gene and intron 11 of the RET gene were found using BLASTN. To avoid interference with transcription, only lncRNAs complementary to the coding strand of intron 1 of the CCDC6 gene were selected. For intron 11 of RET, lncRNAs complementary to both DNA strands were considered because this gene is expressed at a much lower level in the thyroid. The selected lncRNAs had calculated melting temperature of the expected binding regions exceeding 58°C ^28^. In the case of multiple complementary lncRNA regions to intron sequences, the one closest to the most thermodynamically stable homology region of the fused genes was chosen as the putative binding region (Figure S1). A search for lncRNAs potentially capable of inducing PML-RARA, CLDN18-ARHGAP26, TMPRSS2-ERG, SS18-SSX1 and SS18-SSX2 chromosomal rearrangements was performed among non-coding transcripts from the GENCODE database Release 39 ^29^.

### Cloning of lncRNAs

At the first stage, primers without restriction sites were used, and as a template for PCR we selected human genomic DNA (if the lncRNA under study does not contain introns), cDNA from human brain tissue (hippocampus), and mix cDNA from several human cell lines: Nthy-ori 3-1, A549, HEK293T, HepG2, RKO, and MCF7. The PCR template was chosen based on the expression level of the corresponding lncRNAs according to GTEx and ZENBU resources. The resulting products were verified by Sanger sequencing (EIMB RAS “Genome” sequencing center). At the second stage, amplification was performed with primers containing sites of the corresponding restriction enzymes (sequences of primers used are given in Table S1). The obtained lncRNAs were inserted into the expression vector pcDNA3.1/Hygro(+) (Invitrogen, USA) containing the ampicillin resistance gene. The lncRNA sequence of CASTL1 was also cloned into the lentiviral vector LeGO iPuro2 (Lentiviral Gene Ontology Vectors, Germany) containing the puromycin resistance gene.

### Transfection and DMS treatment

Transfection was performed using electroporation with the Neon Transfection System (Life Technologies, USA) by two 20 ms and 980 V impulses in 100 µl tips designed for this instrument. To investigate the effect of candidate lncRNAs on the frequency of CCDC6-RET chromosome rearrangement formation when treated with dimethyl sulfate, 5 million cells were electroporated with 5 µg of plasmids encoding tested lncRNAs or with of the pEGFP-N3 reporter vector (Clontech) as previously described ^30^. The GFP-expressing construct was used to evaluate the efficacy of cell electroporation and the effect of transfection on survival. Transfection with an efficiency of 5% or more was considered successful. 2.5 million electroporated cells were placed in the wells of a six-well plate with 2 ml of medium. After 24 hours, dimethyl sulfate (DMS, Sigma) was added to the medium at concentrations of 6 or 24 mM. After 15 minutes of incubation with DMS at 37 degrees Celsius and 5% CO2, cells were supplemented with a fresh medium that was later replaced twice a week. When cells reached near confluence they were collected for subsequent extraction of total RNA and further analysis.

### RNA isolation and cDNA synthesis

Total RNA was isolated from cells using an ExtractRNA kit (Eurogen, Russia). From 0.5 to 2 million cells were used in each experiment. To eliminate genomic DNA, the RNA was treated with DNase I (Thermo Scientific) followed by its deactivation according to the manufacturer’s protocol. Reverse transcription was performed using the MMLV RT kit (Eurogen, Russia) according to the manufacturer’s protocol. For the cDNA synthesis 4 µg of total RNA were taken with random primers and oligo dT in equal amounts, the total reaction volume was 40 µL.

### Quantitative PCR

Real-time PCR was used to analyze the expression level of the chimeric oncogene CCDC6-RET and lncRNA CASTL1. The study was performed on an Applied Biosystems 7500 Real-Time PCR. The reagent mixture qPCRmix-HS LOWROX (Eurogen, Russia) was used for Real-Time PCR with TaqMan probe, which binds to the junction site of two exons of different genes, thus characterizing the expression level of the hybrid RNA. qPCRmix-HS SYBR+LOWROX was used for the evaluation of reference gene and lncRNA levels. The sequences of primers and TaqMan probe used are given in Table S1. Each reaction was performed in 25 µL containing 0.7 µL of cDNA mix when analyzing the RNA level of the reference gene and lncRNA or 3.3µL when analyzing the expression of the fusion oncogene. The TaqMan probe was used at a final concentration of 300 nM, and primers were taken at a concentration of 200 nM. All samples were amplified in three replicates, and the mean was used for further analysis.

### Irradiation

To investigate the ability of CASTL1 lncRNA to affect the frequency of formation of the inversion under irradiation-assisted double-strand breaks conditions, 5 million cells were transfected with 5 µg of pcDNA-3.1-CASTL1 plasmid or pcDNA-3.1 control vector. The cells in the standard medium were placed in T12.5 Tissue culture treated flasks with filtered caps and transported to the N.N. Blokhin Research Center for Oncology, Ministry of Health of Russia, where the cells were exposed to 6 MV bremsstrahlung radiation at the SL75-5-MT linear electron accelerator (distance to the emitter 100 cm, field 30 × 30 cm), where the plates with cells were placed between the 2 cm thick Plexiglas plates to establish primary electrons equilibrium. The control flask was transported but not irradiated. Doses of 1, 4, 9, 16, and 25 Gy were used. 1 Gy did not cause any effect while 25 Gy resulted in massive cell death. Therefore, doses of 4, 9 and 16 Gy were used in the experiments.

### CRISPR/Cas9 and cell sorting

The crRNAs targeting 1 and 11 introns of the CCDC6 and RET genes respectively were designed using E-CRISP (e-crisp.org) ^31^ and cloned into pSpCas9(BB)-2A-GFP (PX458) construct which was a gift from Feng Zhang (Addgene plasmid #48138) ^32^. 5 µg of lncRNA-carrying expression vector was co-transfected together with 7.5 µg of each of the PX458-derived constructs. The cells were placed in 2 wells of a 6-well plate, 2.5 million each, in 2 ml of complete medium. Twenty-four hours after electroporation, 30 000 to 60 000 GFP-positive cells were selected using Fluorescence-activated cell sorting (FACS) on the S3e Cell Sorter (Bio-Rad Laboratories, USA). 1 mL of RPMI-1640 medium was added to the receiving tube, medium with 5nM Na2Se03 was used in some experiments to reduce the effect of oxidative stress on DNA damage and improve overall cell survival with CRISPR/Cas after sorting ^33^.After sorting, cells were placed in 12-well plates and grown on Na2Se03 medium for 3 days. Subsequently, medium without selenite was used to maintain the cells. Once 80-90% confluence was achieved, cells were used to isolate total RNA.

### Statistical analysis

Statistical analysis was performed using Graphpad Prism 9 software. A one-sample Wilcoxon test was applied to the data normalized to the control sample, unless otherwise stated in the figure description. Absolute values were compared using the Mann-Whitney test. Spearman correlation analysis was performed using the Python package SciPy.

### Analysis of RNAseq data from tumors and normal thyroid tissues

Data from the TCGA REBC-THYR project ^18^ was used to analyze CASTL1 expression in tumors and normal thyroid tissues. This project contains the largest collection of tumor and normal thyroid tissues, with extensive coverage of RNA samples. This is important in the study of CASTL1 expression because most tumors and normal tissues have this RNA at a low level in bulk tissue. The Mann-Whitney test was used to compare expression levels. To find genes whose expression changes with CASTL1 expression in normal thyroid tissues, the Spearman correlation was calculated. The 2000 genes whose expression correlated with CASTL1 level with the lowest p-values were used for gene group enrichment analysis using the Metascape service ^34^. Among noncoding RNAs without known alternative transcripts and with expression in thyroid similar to CASTL1 (median and interquartile range ratios less or equal 1.1) three genes were randomly chosen as controls for correlational analysis.

## Results

### 1. Search for lncRNAs potentially capable of stimulating CCDC6-RET

CCDC6-RET inversion results in joining intron 1 of CCDC6 gene and intron 11 of RET gene ^15^ in the vast majority of cases, so sequences of these introns were used to identify four lncRNAs from GENCODE database with putative binding regions having an estimated melting point at 58°C or higher (Table S2). All sequences of candidate lncRNAs that potentially bind to intron 1 of CCDC6 gene are SINE repeats of AluS family, whereas one of the sequences potentially able to bind to intron 11 of RET gene from CASTL1 lncRNA is LTR repeat sites of ERV1 family. This is consistent with our suggestion that RE sequences have the potential to facilitate interaction between lncRNAs and introns of protooncogenes.

### 2. CASTL1 stimulates CCDC6-RET formation upon DNA damage by dimethyl sulfate

Previous studies ^4^ suggested that RNAs capable of interacting with sites of two genes can promote their fusion during the reparation of double-strand DNA breaks. For an initial assessment of the ability of lncRNAs to induce the formation of the desired inversion, an alkylating agent dimethyl sulfate (DMS) was used to create a large number of random DNA breaks ^35,36^. In this model, we examined the ability of CASTL1, CYP4F8-006 and AC068282.3-001 lncRNA transcripts to increase the frequency of CCDC6-RET inversion. We found that lncRNA CASTL1, but neither CYP4F8-006 nor AC068282.3-001, significantly increased the rate of CCDC6-RET chimeric transcript formation when cells were treated with DMS. (Figure 1A). Sequencing of the CCDC6-RET amplicon showed that it indeed contained the spliced sequences of exon 1 of the CCDC6 gene and exon 12 of the RET gene (Figure 1B). Based on these data, CASTL1 lncRNA was selected to further test the effect in other models.

**Figure 1:**
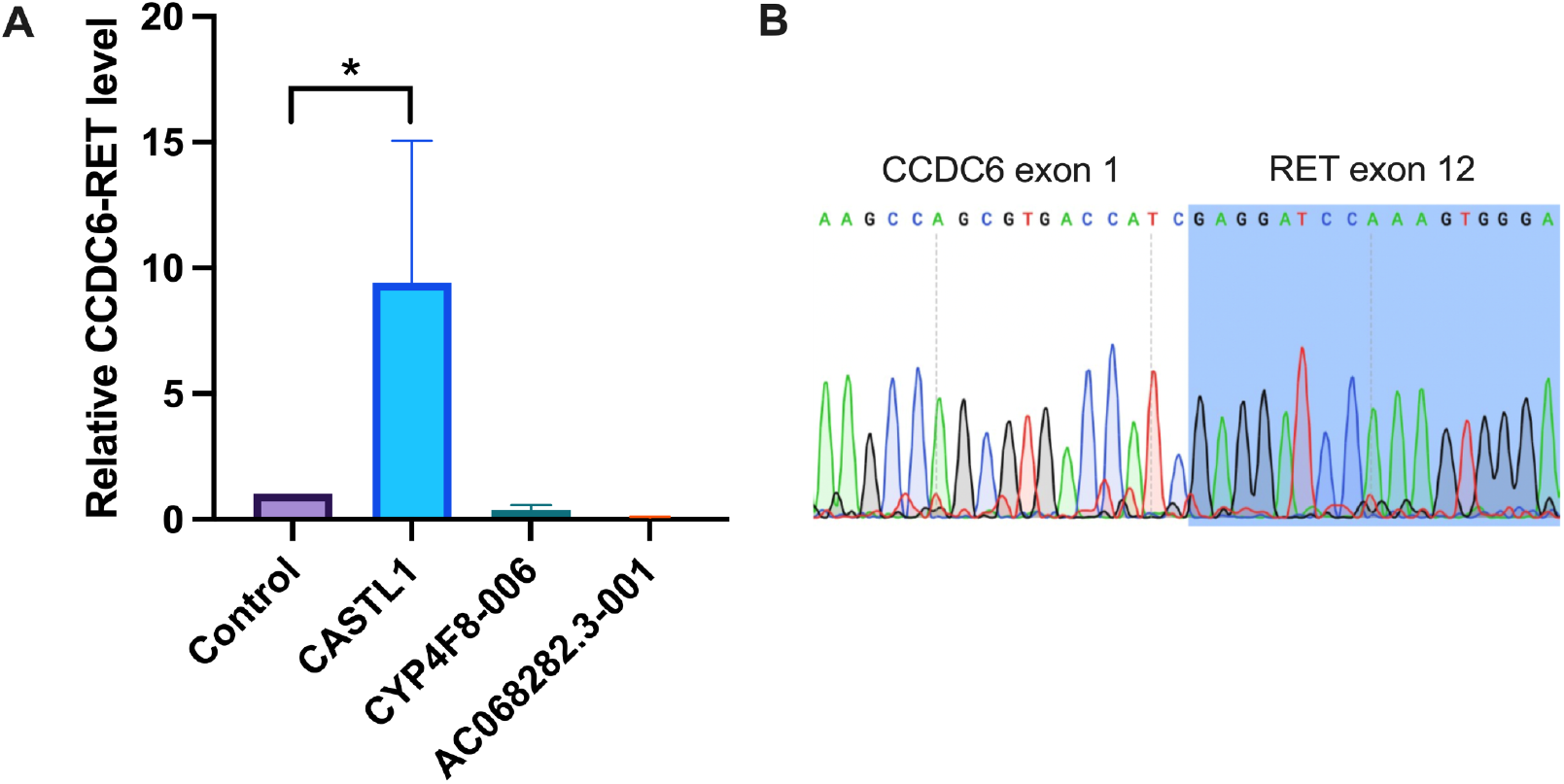
Assessment of the ability of CASTL1, CYP4F8-006 and AC068282.3-001 lncRNAs to induce CCDC6-RET chromosomal rearrangement. A. Expression level of CCDC6-RET in Nthy-ori 3-1 cells transfected with the indicated lncRNAs and treated with 24 mM DMS. The expression of the chimeric oncogene was normalized to the expression of beta-actin. Control samples of Nthy-ori 3-1 (GFP) cells were transfected with pE-GFP-N3 plasmid. Means ± SEM from 3 independent experiments are presented. *denotes p-values of less than 0.05. B. Sequence of CCDC6-RET junction in the chimeric PCR product.

**Figure 2:**
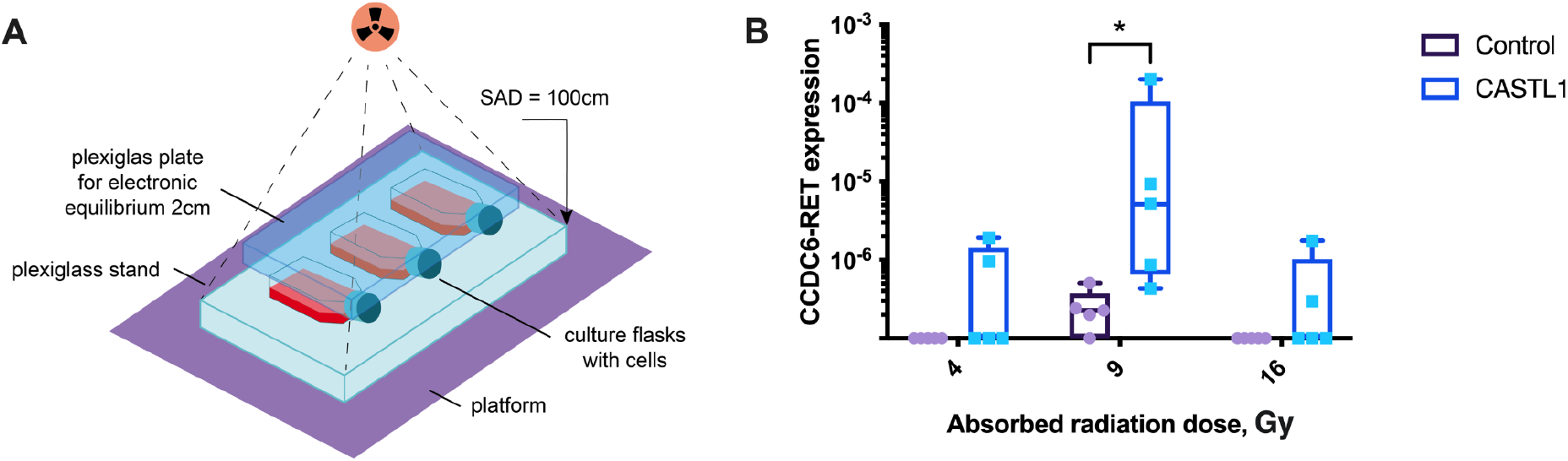
Assessment of the ability of CASTL1 lncRNA to stimulate CCDC6-RET chromosomal rearrangement under irradiation conditions. **A**. Layout of the irradiation experiments. **B**. CCDC6-RET chimeric oncogene expression was normalized to beta actin. Control samples of Nthy-ori 3-1 cells were transfected with the original pcDNA-3.1 vector, samples of CASTL1 cells were transfected with pcDNA-3.1-CASTL1 plasmid. Data from 5 independent experiments are presented. The horizontal bar indicates the median. The data on the X axis indicate any results not exceeding the level of negative control (amplification without template). * denotes a p-value less than 0.05 according to the Mann-Whitney test.

### 3. CASTL1 increases the probability of CCDC6-RET inversion upon irradiation

In order to test whether CASTL1 lncRNA can induce CCDC6-RET inversion in Nthy-ori 3.1 cells on its own, we expressed this RNA from a lentiviral vector but did not observe a significant level of the chimeric oncogene CCDC6-RET even after six months, presumably due to an insufficient level of double-strand breaks. This is consistent with many patients with CCDC6-RET aberration having a history of radiation exposure ^37,38^. Therefore, we hypothesized that the use of radiation to introduce DNA breaks would better correspond to the conditions that favor the chromosomal aberration. Under irradiation with a dose of 9 Gy the highest level of chimeric oncogene expression was observed in cells expressing CASTL1 lncRNA while at doses of 4 and 16 Gy it was relatively low. Presumably, at the dose of 4 Gy there are insufficient DNA breaks while at the dose of 16 Gy the irradiated cells are less likely to survive. This nonlinear relationship echoes the observation that patients who received different doses of radiation are characterized by different chromosomal rearrangements ^39^.

### 4. CASTL1 stimulates CCDC6-RET fusion when targeted DNA breaks are introduced by CRISPR/Cas9 complexes

Irradiation and DMS introduce non-directional DNA damage that limits the maximum intensity of treatment due to excessive cell death. Therefore, further studies on the effect of lncRNAs on the CCDC6-RET inversion were performed using endonuclease CRISPR/Cas9 complexes.

In pilot experiments when all GFP-positive cells were selected after transfection, the level of CCDC6-RET chimeric gene transcripts was independent of CASTL1 expression. This might be due to the excessive amount of double-strand breaks in the introns of CCDC6 and RET genes introduced by the CRISPR/Cas9 complexes that itself was sufficient to initiate chromosomal rearrangements ^40^. In order to use cells with lower levels of CRISPR/Cas9 complexes, we divided the GFP-positive cell population into two approximately equal subpopulations of low and high fluorescent cells (Supplementary Material FIG. S2). In this experiment, the level of CCDC6-RET fusions in cells expressing CASTL1 was significantly different from the control sample in the low fluorescent subpopulation (FIG. 3A). At the same time, in the high fluorescent subpopulation, the expression level of the chimeric oncogene was independent of the CASTL1 expression, which confirmed our hypothesis (FIG. 3B).

**Figure 3:**
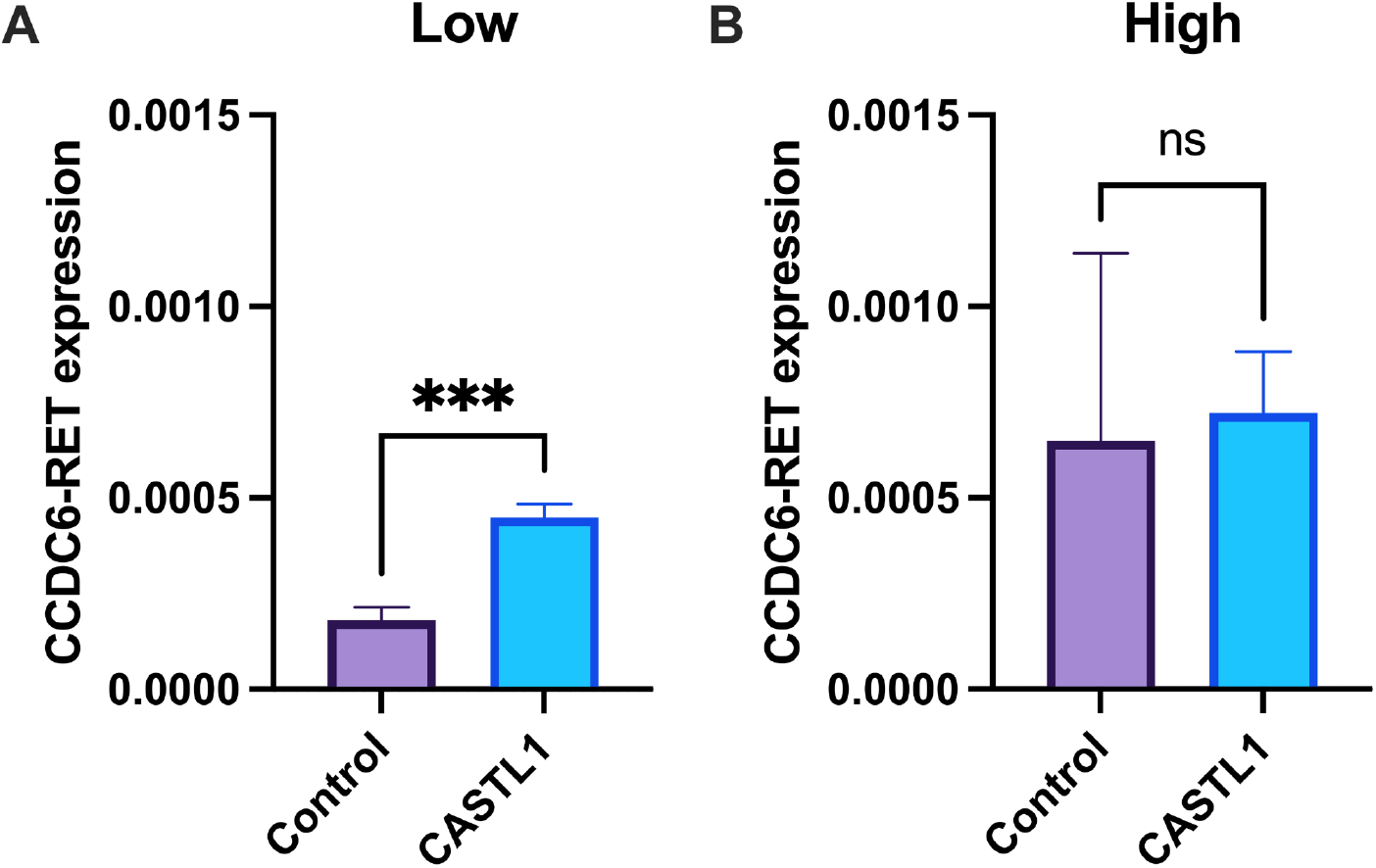
CASTL1 increases the incidence of CCDC6-RET in Nthy-ori 3-1 cells with low activity of CRISPR-Cas9 endonuclease complexes. **A, B**. CCDC6-RET levels in control of Nthy-ori 3-1 cells and cells expressing CASTL1 in subpopulations with low (A) or high (B) levels of CRISPR-Cas9 expression. Chimeric oncogene expression was normalized to beta actin. Control samples are Nthy-ori 3-1 cells transfected with an empty vector. Means ± SD for 3 independent experiments are shown. ***means p-value less than 0.001 by Student’s test, ns-means no significant difference between mean CCDC6-RET expression levels.

### 5. Regions of homology to CCDC6 and RET genes are required for CASTL1 ability to stimulate CCDC6-RET

In previous experiments we observed that cells expressing CRISPR/Cas complexes often did not survive well after sorting. To improve the viability of transfected cells, Na2Se03 was added to the culture medium to reduce oxidative stress. Under these conditions, the effect of CASTL1 lncRNA became more pronounced (Figure 4), allowing a reliable measurement of a decline in lncRNA activity due to the deletion of a particular RNA sequence.

**Figure 4:**
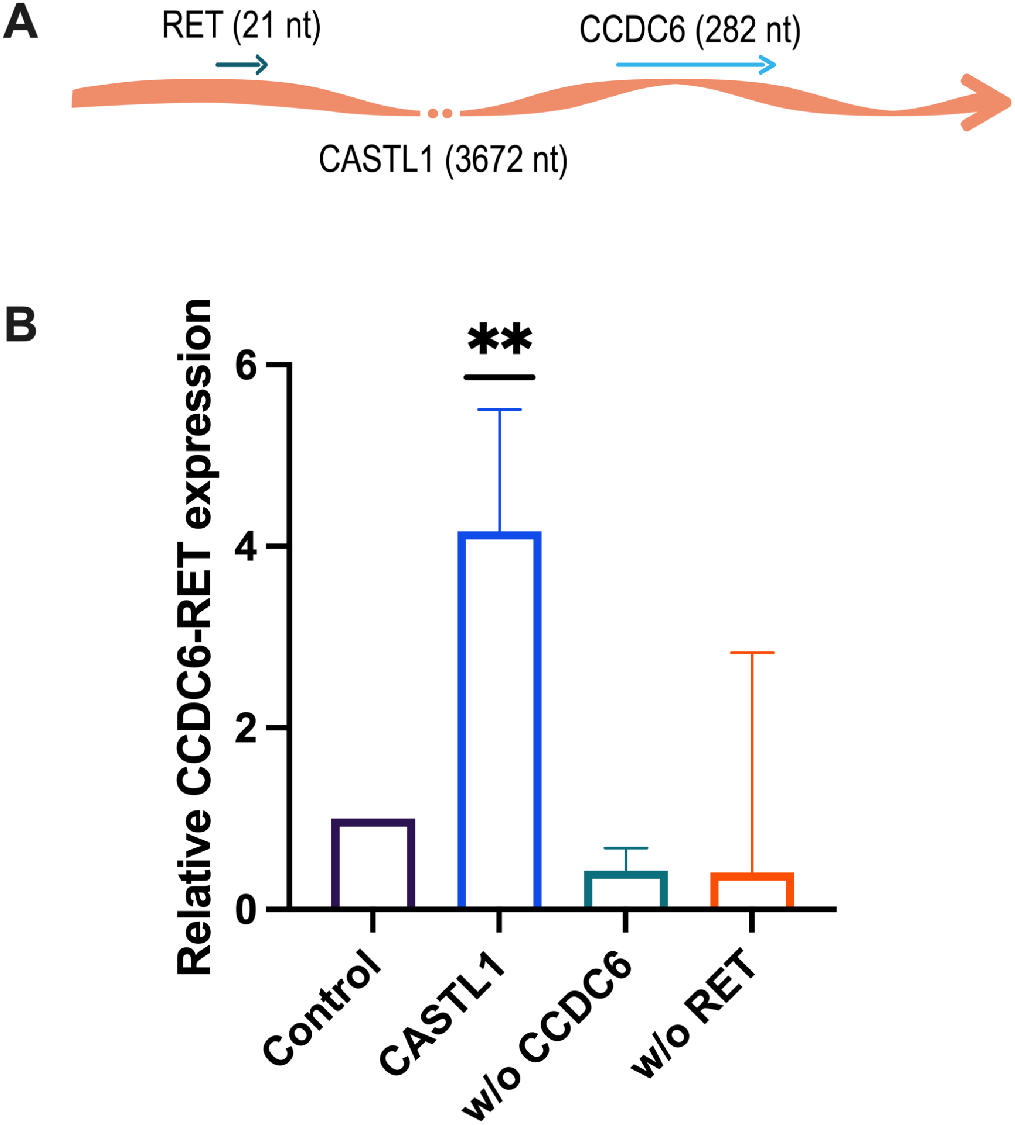
CASTL1 lncRNA but not its deletion variants can stimulate CCDC6-RET inversion in Nthy-ori 3-1 cells. **A**. Schematic of the location of the putative interaction sites of CASTL1 with the CCDC6 and RET gene sequences **B**. The expression of chimeric oncogene CCDC6-RET was normalized to beta-actin gene and then to the values obtained for the control sample transfected with empty vector pcDNA-3.1. The abscissa axis shows the examined cell samples transfected with plasmids carrying the sequence of different lncRNA variants RP6-91H8.5 including the initial variant CASTL1 and its derivatives lacking the putative interaction regions with intron 1 sequence of CCDC6 (w/o CCDC6) and intron 11 of RET gene (w/o RET). Median and interquartile ranges from 8 independent experiments are presented. **-means the difference from control sample with p-value less than 0.01 by Wilcoxon test

To show the functional significance of the regions of CASTL1 homology to introns of CCDC6 and RET genes, we constructed mutant versions of CASTL1 without these regions. Our data showed that both CASTL1 deletion variants did not significantly increase the expression of the chimeric CCDC6-RET oncogene, demonstrating the importance of these regions in the rearrangement process. Interestingly, the median expression level of CCDC6-RET for the CASTL1 deletion variants was slightly lower than that of the control, probably because CASTL1 binding to only one of the introns inhibited the formation of this rearrangement or directed the formation of other chromosomal rearrangements.

### 6. Thyroid tumors with CCDC6-RET translocation show increased expression of CASTL1

If CASTL1 expression facilitates CCDC6-RET fusion, the expression of this lncRNA can be expected to remain elevated in the cells where this fusion has already occurred, especially in highly differentiated tumors where malignant cells retain characteristics of the original cells from which they develop. To this end, we analyzed papillary thyroid carcinoma, a tumor characterized by high degree of differentiation ^41^, using RNAseq data from the REBC-THYR (Comprehensive genomic characterization of radiation-related papillary thyroid cancer in Ukraine) project ^18^. The analysis showed that CASTL1 levels were indeed significantly higher in tumors with CCDC6-RET inversion (Figure 5A). CASTL1 levels in normal thyroid tissues demonstrated a weak (11%), significantly indistinguishable increase in median CASTL1 levels in samples from patients with tumors carrying the CCDC6-RET mutation (Figure 5B). Based on this, we hypothesize that the elevated level of CASTL1 expression in CCDC6-RET tumors may reflect its increased expression and possible involvement in CCDC6-RET translocation in the original cells.

**Figure 5.**
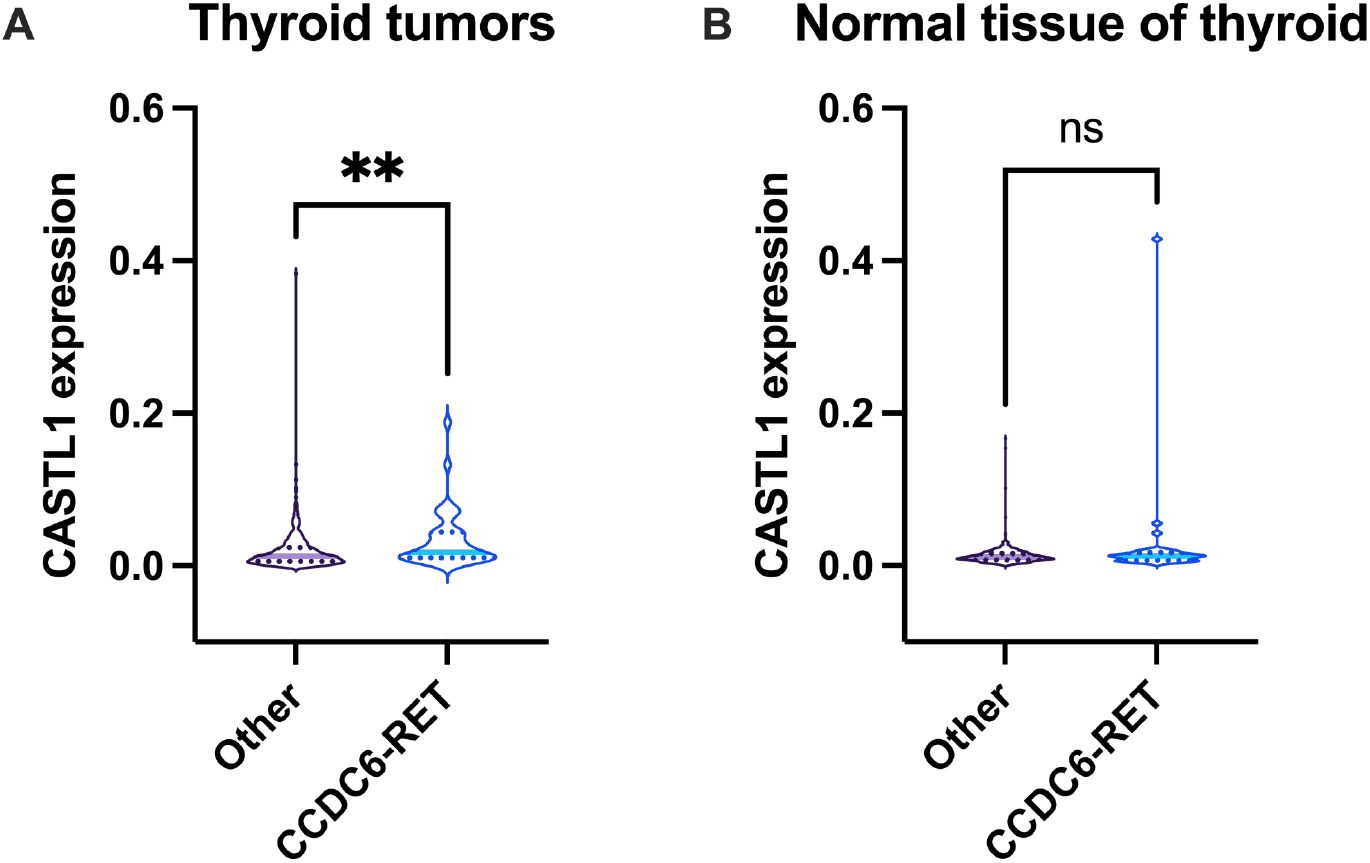
Comparison of CASTL1 lncRNA expression in thyroid tumors with CCDC6-RET rearrangement and in tumors with other driver mutations. The level of CASTL1 is shown in FPKM (Fragments per kilobase of transcript per million mapped fragments). **A**. CASTL1 expression in papillary thyroid carcinoma cells. **B**. CASTL1 expression in normal thyroid tissue from patients with papillary carcinoma tumors.** denotes the difference with the control sample with p-value less than 0.01 by Mann-Whitney test.

### 7. Expression of CASTL1 in normal thyroid tissue correlates with the expression of genes associated with autoimmune and hematological diseases

To estimate possible physiological causes of increased CASTL1 expression in normal thyroid tissue, we performed a correlation analysis of CASTL1 and other genes co-expression. It revealed that CASTL1 is positively correlated with the expression of a number of genes related to immune cell activity (Figure 6A). Analysis of the association of CASTL1 levels with disease-specific transcriptome changes revealed correlation with the expression of genes involved in autoimmune diseases, leukemias, lymphomas and immunodeficiencies (Figure 6B). Control genes with similar expression levels in the thyroid showed entirely different patterns (Figure S3), indicating that association with immune responses was specific to CASTL1. The list of transcription factors whose activity may correlate with CASTL1 expression generated using the TRRUST resource ^42^ was clearly enriched in the factors involved in inflammatory reactions (Figure 6C). According to the GTRD collection of uniformly processed ChIP-seq data (gtrd.biouml.org/#) ^43^, most top hits from this list (SPI1, RELA, TP53, EP300) can interact with the CASTL1 gene locus (Figure S4), providing a mechanistic basis for their possible involvement in the regulation of CASTL1 expression in thyroid tumors. Overall, our analysis suggests a possible relationship between CASTL1 expression and inflammatory response. This suggestion is consistent with the detection of the CCDC6-RET driver chromosomal rearrangement in patients with autoimmune thyroid disease even before tumor development ^44^.

**Figure 6:**
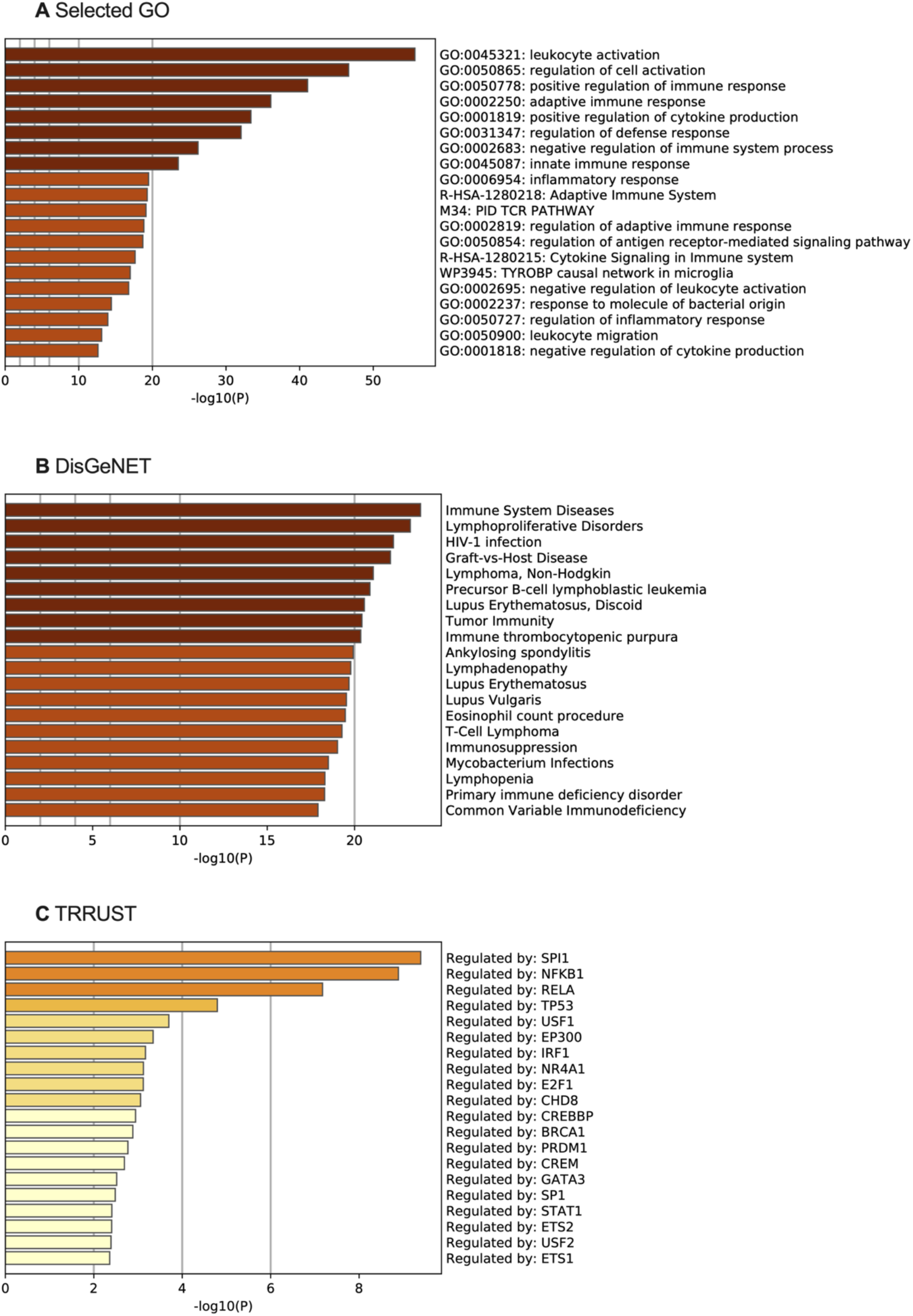
Analysis of genes whose expression correlates with CASTL1 lncRNA levels in thyroid cells. A. Enrichment analysis of functional groups of genes whose expression correlates with CASTL1. B. Identification of diseases characterized by transcriptome changes correlating with increased levels of CASTL1. C. Identification of transcription factors whose activity is associated with changes in gene expression that correlate with an increase in CASTL1 levels.

## Discussion

By now RNA molecules have been established as key players in most genome related processes ^13,45,46^. However, only a few works have been devoted to the role of RNA in the stimulation of directed chromosome rearrangements in vertebrates. In 2008 the “the cart before the horse” hypothesis was proposed based on the detection of chimeric JAZF1-JJAZ1 mRNA in normal endometrial cells without corresponding oncogenic fusion ^47^. The hypothesis stated that chimeric mRNA formed as a result of trans-splicing can stimulate the fusion ^48^. More recent studies have shown that trans splicing products are mostly detected at very low levels in human cells ^49^. Moreover, some papers suggest that such RNAs can appear as a result of template switching during reverse transcription in vitro ^50^. These observations cast doubt on the contribution of such chimeric RNAs to mutagenesis. In addition, “sense” chimeric RNA, which mimics a possible trans splicing product, stimulates the formation of aberration only when RNA polymerase II activity is inhibited by a-amanitin ^4,51^.

In this study, we demonstrated that lncRNA CASTL1, containing regions complementary to the intron sequences of CCDC6 and RET, can stimulate CCDC6-RET chromosomal rearrangement. Analysis of recently published data showed that tumors with this translocation are indeed characterized by increased levels of this lncRNA. To assess the prospects of generalizing the phenomenon of RNA-facilitated chromosomal rearrangements, we searched for lncRNAs and mRNAs potentially capable of causing other driver oncogenic rearrangements, limiting the search to frequent gene fusions (more than 4% of cases for tumor types from TCGA projects): PML-RARA frequently found in Acute Myeloid Leukemia samples ^52^, CLDN18-ARHGAP26 characteristic of Gastric Adenocarcinoma ^53^, TMPRSS2-ERG which is the main driver mutation of Prostate Adenocarcinoma ^54^, and SS18-SSX1 and SS18-SSX2 found in the vast majority of Synovial sarcoma ^55^. The fusion regions of these genes were taken from ChimerKB4 ^3^. In case of longer introns, the lists of RNAs potentially capable of driving such gene fusions contain hundreds or even thousands of transcripts Most of the abovementioned frequent rearrangements have large fused introns which make lists of candidate RNAs quite long, however in the case of CLDN18-ARHGAP26 fusion only 203 candidate transcripts were found Table S3 (in a separate file). Candidate lists can be further reduced by filtering RNAs by length, complementarity to the coding strand, and expression in the relevant cell type. It is worth noting that the regions of potential interaction of many lncRNAs with the introns of the genes under study are often RE sequences. Although these sequences provide large regions of potential interaction between lncRNAs and the introns, they may prevent lncRNAs from stimulating the formation of certain chromosomal rearrangements. First of all, multiple RE copies in the genome will compete for interactions with the same lncRNA. Furthermore, adenines in the RE sequences within the lncRNA can be modified by enzymes of the adenosine deaminase RNA specific (ADAR) family while interacting with a complementary sequence of the same or another RNA molecule ^56^. The secondary structure of lncRNAs or peculiarities of their localization could also prevent binding to complementary DNA. Further discovery and study of RNAs capable of stimulating chromosomal rearrangements may reveal additional search criteria to further shorten the list of candidate RNAs.

Analysis of the expression profile of normal thyroid tissue samples showed that the level of CASTL1 RNA correlates with the expression of genes involved in inflammatory reactions, primarily in autoimmune processes. At the same time, chronic inflammation associated with autoimmunity or radiation damage is known to increase the likelihood of developing thyroid cancer ^44,57^. The inflammation promotes tumorigenesis via reactive oxygen species that cause mutations, via regenerative growth following cell death in normal tissue as well as by pro-inflammatory cytokines that contribute to the release of insulin and thyrotropic hormones that stimulate tumor cell growth. Moreover, immune cells involved in chronic inflammation, especially macrophages, contribute to tissue reorganization and vascularization necessary for tumor growth ^57–59^. Our data reveal another possible factor of oncogenesis in thyroid associated with inflammation: increased levels of CASTL1 lncRNA which stimulates CCDC6-RET aberration. This suggestion is consistent with a significant association of thyroid autoimmunity with CCDC6-RET but not with the other two most frequent driver mutations characteristic of papillary thyroid carcinoma, BRAFV600E and NCOA4-RET ^60^.

In our work, we demonstrated that the usage of certain RNAs can facilitate targeted genome rearrangements under conditions of random DNA damage and during the expression of CRISPR/Cas9 complexes. The use of RNA interacting with genomic DNA near target sites in combination with CRISPR/Cas9 can be used to increase the efficiency of genome modifications.

Thus, we have shown for the first time that a lncRNA can stimulate a driver chromosomal rearrangement. Suppression of such lncRNAs expression may become a basis for novel therapeutic methods to prevent the formation of primary and secondary tumors, particularly those associated with DNA-damaging factors ^61^. Also, when introducing new CRISPR/Cas9-based therapies, possible side-effects of rearrangement formation should be taken into account and the expression levels of CASTL1 and similar lncRNAs may need to be controlled.

## Acknowledgments

Work on most sections of this article was supported by the Russian Science Foundation (project no. 19-74-10083). Search for potentially oncogenic RNAs for the discussion section supported by grant 075-15-2019-1660 from the Ministry of Science and Higher Education of the Russian Federation. We would like to thank Grigory Efimov from the National Research Center for Hematology for providing some reagents. We are thankful to Violetta Gogoleva from Engelhardt Institute of Molecular Biology for her help with cell sorting.

## Author Contributions

Conceptualization, A.M.S., D.E.D., A.N.U. and M.M.M.; investigation, D.E.D., M.M.M., A.M.S, A.N.U., E.Y.S, E.A.T, V.V.K., P.V.S., G.E.G, S.A.S; computational analysis, D.E.D; writing—original draft preparation, M.M.M., A.M.S., K.A.T., A.S.U. and D.E.D.; writing—review and editing, K.V.K., D.E.D.,D.V.K. and M.M.M.; visualization, E.M.S.; funding acquisition, A.M.S. and D.V.K. All authors have read and agreed to the published version of the manuscript.

## Supplementary

**Figure S1.**
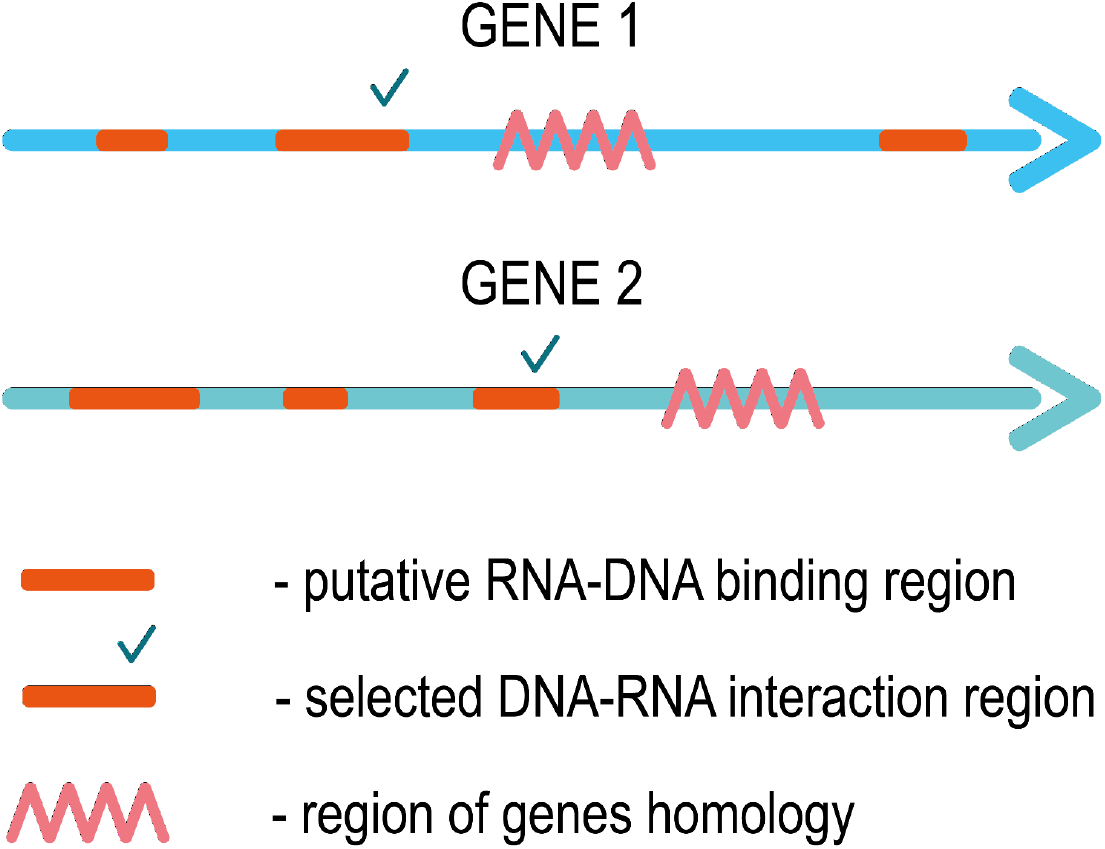
Complementary regions of lncRNA and intron sequences. Selection of the putative lncRNA binding sites closest to the most thermodynamically stable homology region of the introns of the fused genes.

**Figure S2.**
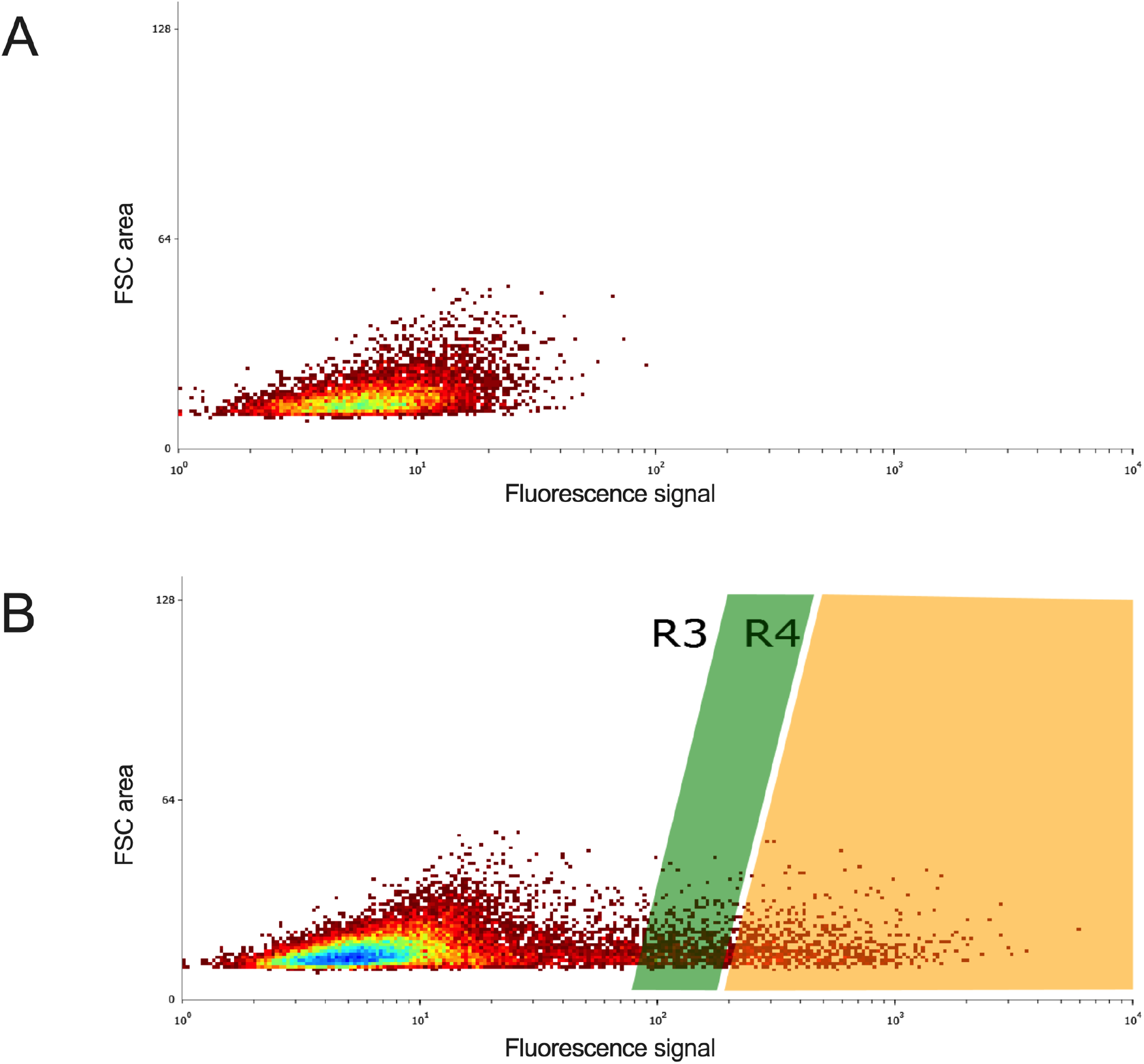
Gates for division of populations of transfected cells. **A**. Level of cell autofluorescence. **B**. Selection of two cell subpopulations by GFP luminescence level: lower fluorescence **low** (R3) and higher fluorescence **high** (R4).

**Figure S3:**
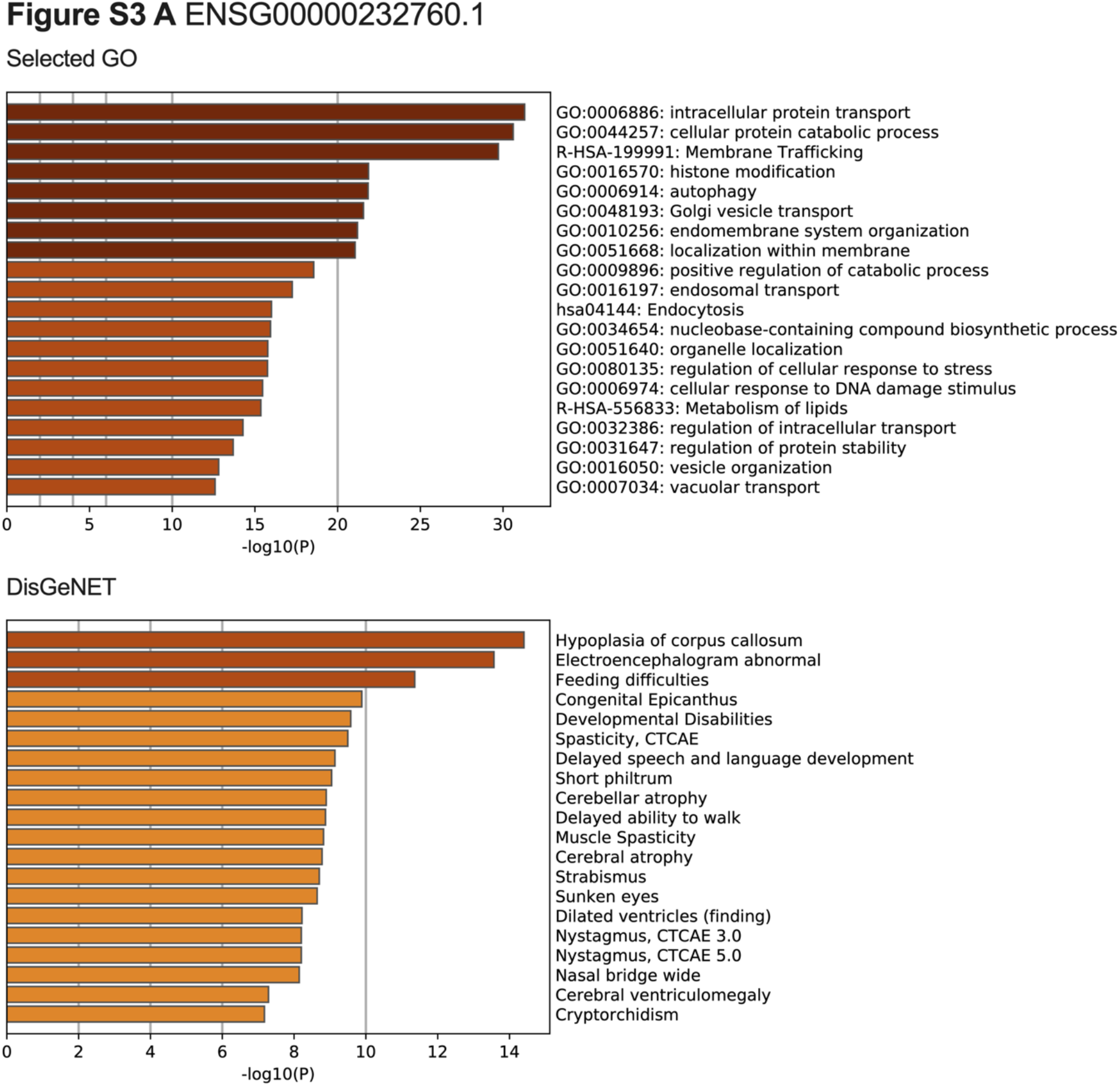
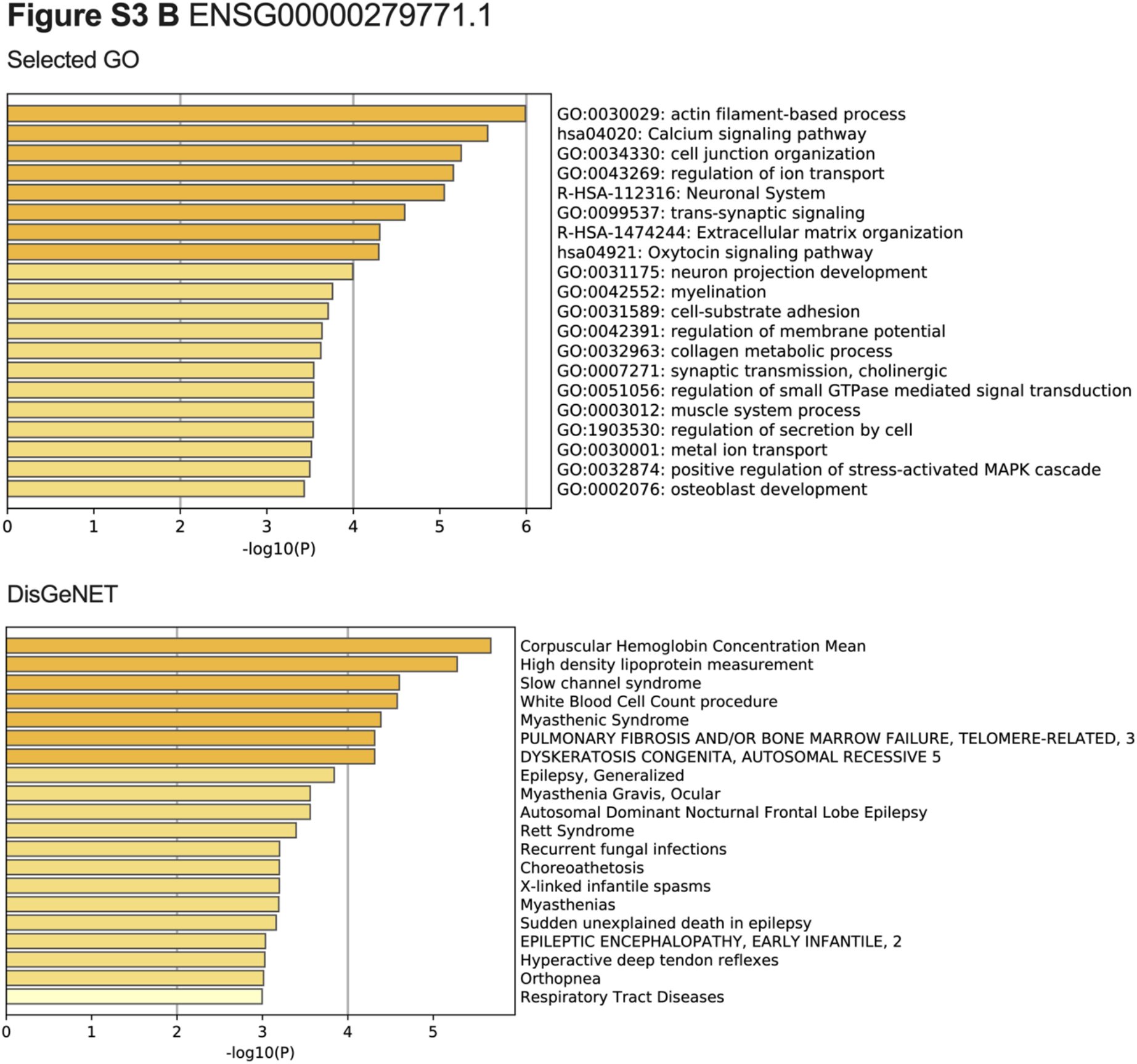
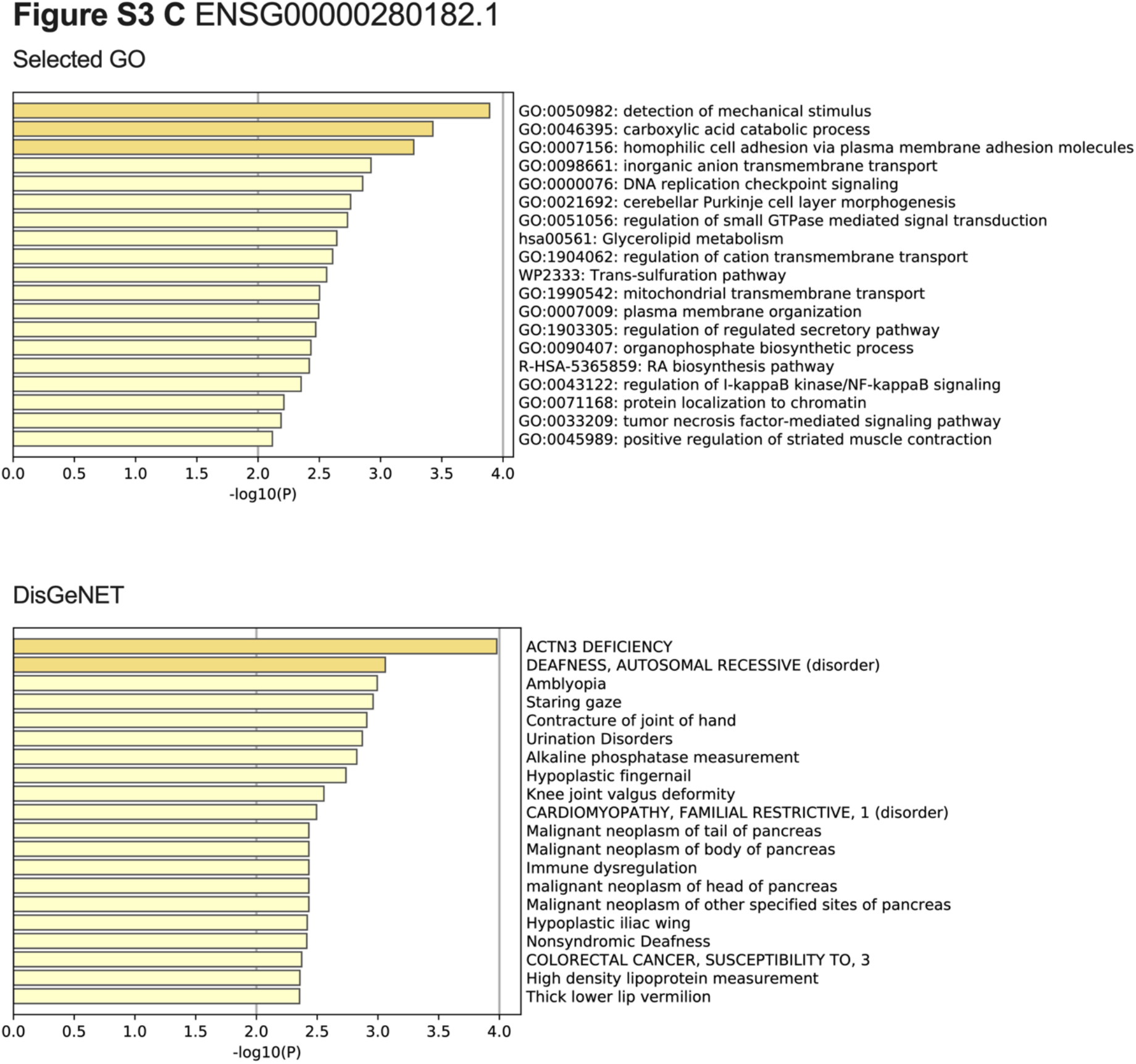
Correlational analysis of control genes. Three genes with expression in thyroid cells similar to CASTL1 were selected for enrichment analysis. Functional groups of genes whose expression correlates with ENSG00000232760.1 (A), ENSG00000279771.1 (B), ENSG00000280182.1 (C).

**Figure S4.**
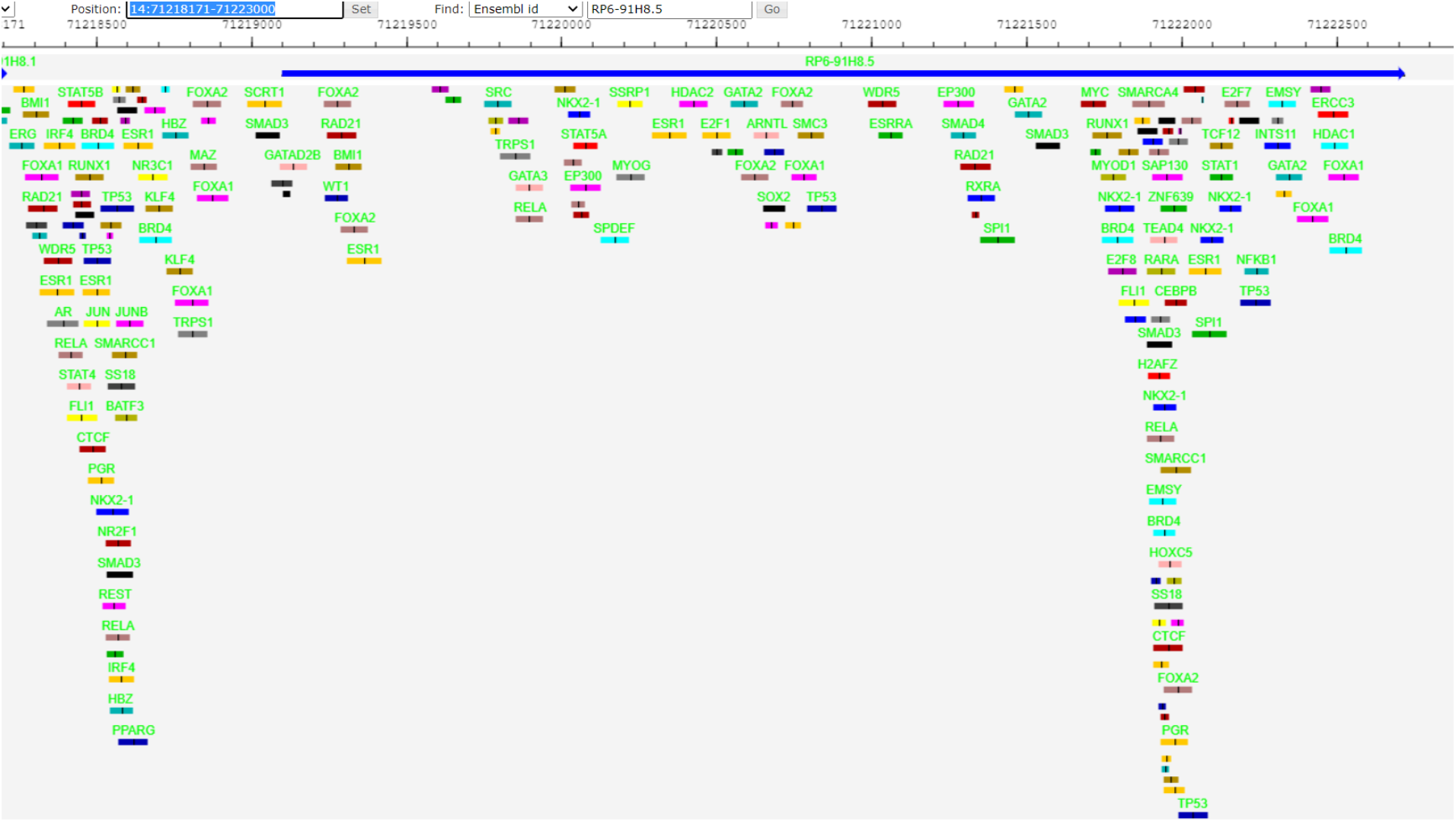
Transcription factor binding regions at the CASTL1 gene locus, according to ChIP-seq data. The blue arrow denotes the CASTL1 sequence; the colored rectangles denote the binding regions of the transcription factors according to CHiP-seq data.

**Table S1.**
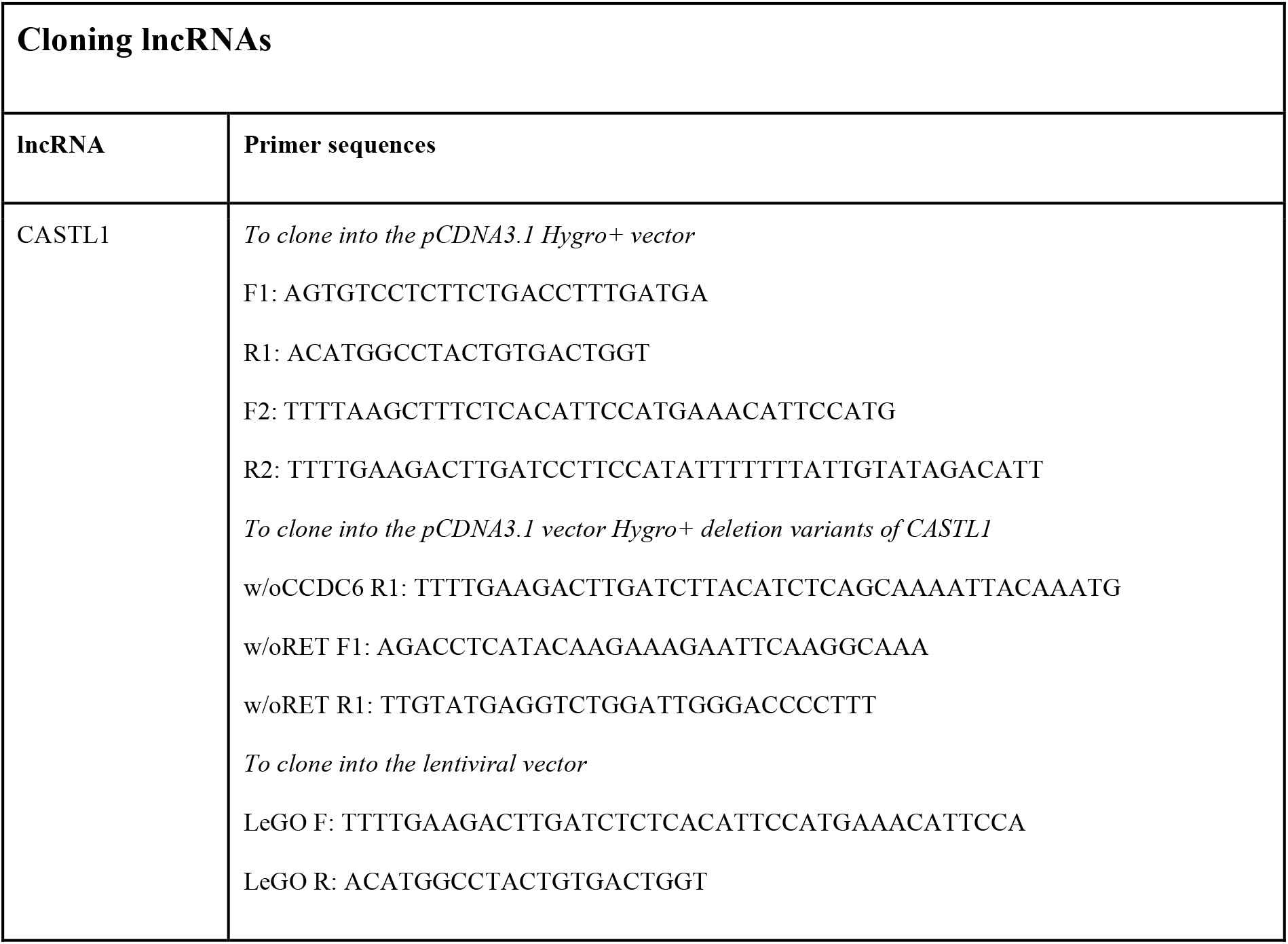
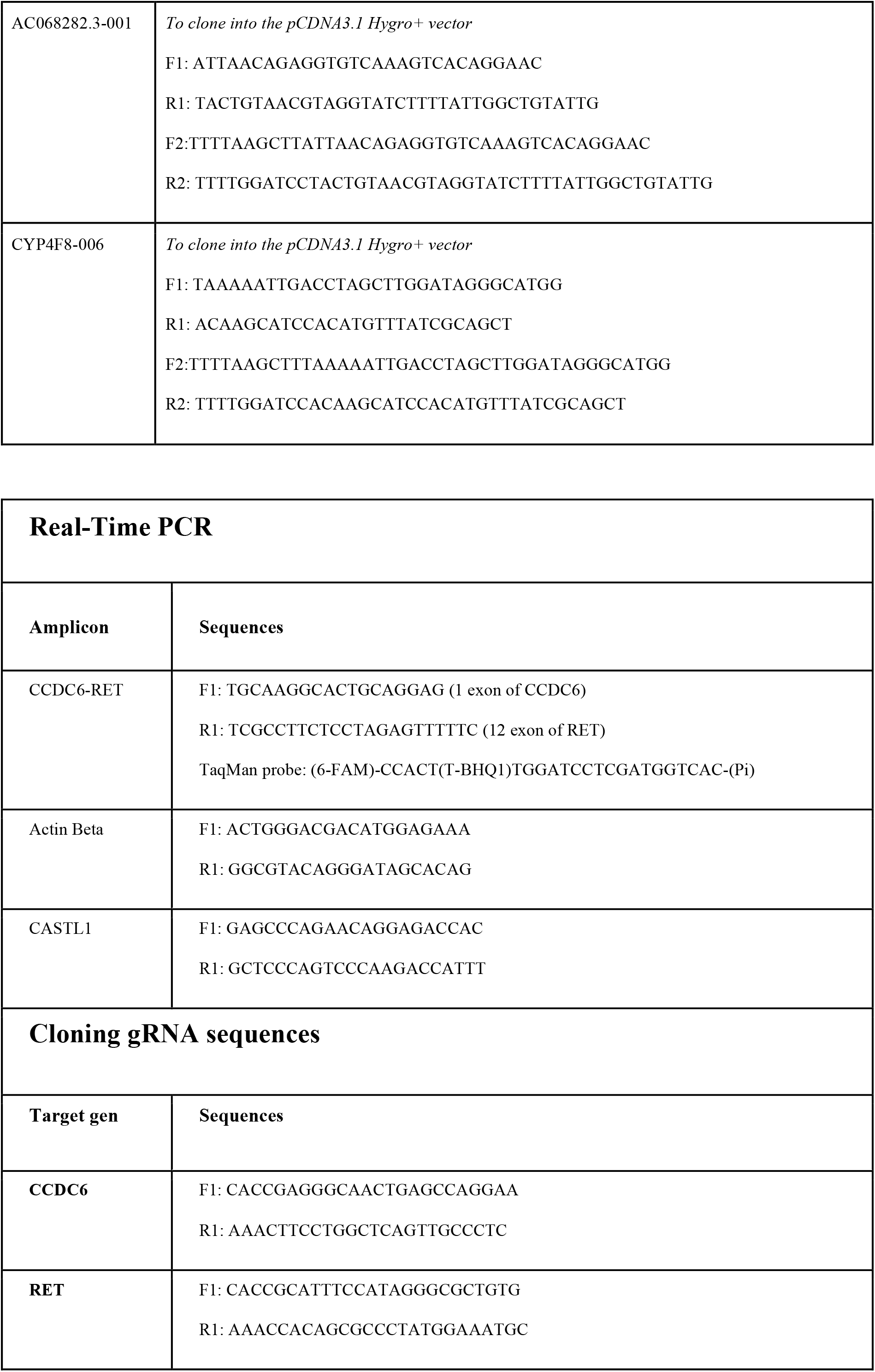
Oligonucleotides.

**Table S2.**
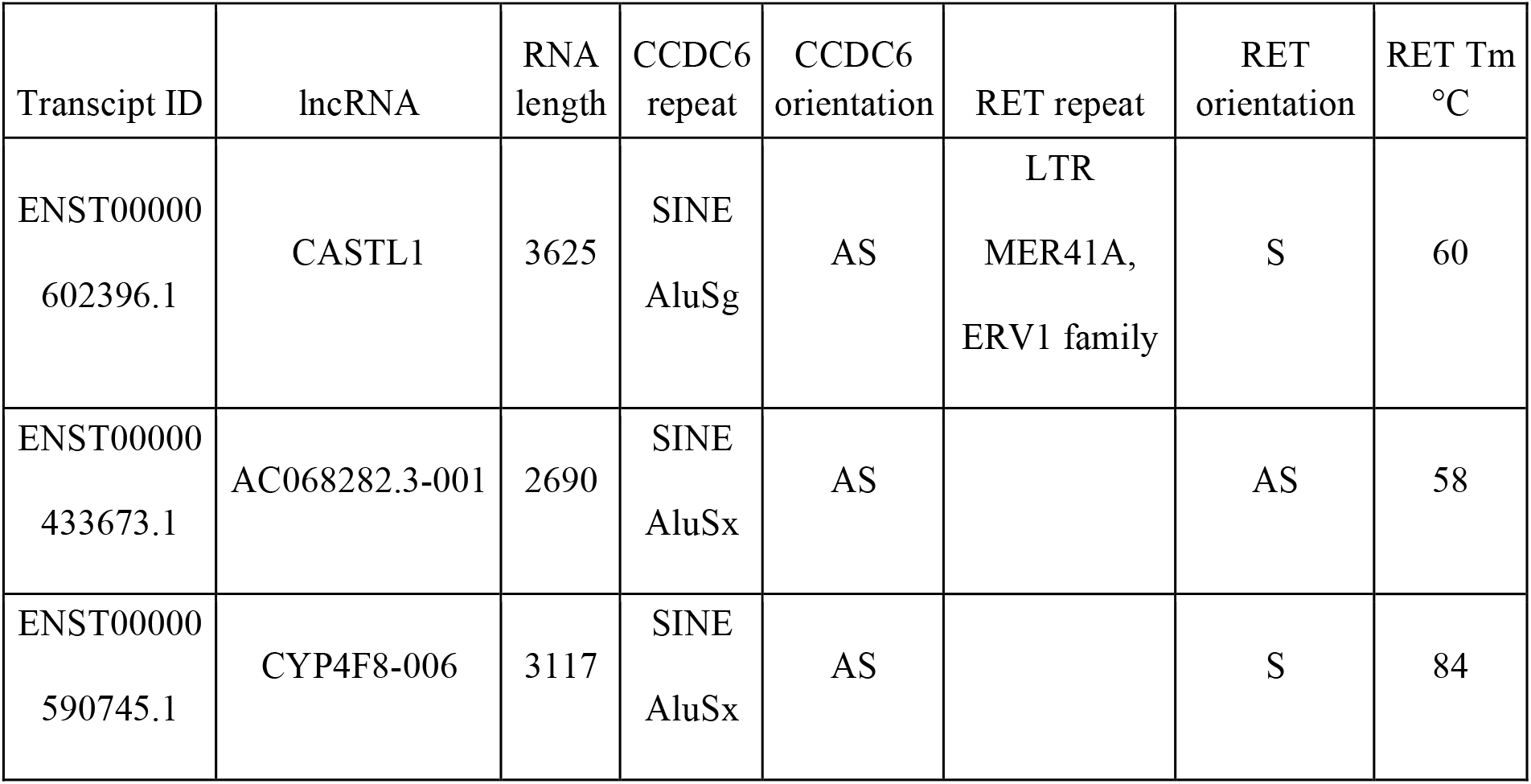
lncRNAs selected for the study. lncRNAs potentially capable of stimulating CCDC6-RET chromosomal rearrangement. The CCDC6 repeat and RET repeat – columns indicate which repetitive element sequence represents the putative interaction region of a given lncRNA with intron of a given gene, CCDC6 repeat and RET repeat orientation – indicates the orientation of lncRNA sequence potentially interacting with intron of a gene relative to its mRNA (S-sense, AS-antisense), Tm °C (RET) – estimated melting temperature of putative lncRNA binding region with 11 intron sequence of RET gene. The Tm of the putative lncRNA interaction region with 1 intron of the CCDC6 gene for all RNAs exceeded 60°C.

| Transcript ID | lncRNA | RNA length | CCDC6 repeat | CCDC6 orientation | RET repeat | RET orientation | RET T<sub>m</sub> °C |
| --- | --- | --- | --- | --- | --- | --- | --- |
| ENST00000602396.1 | CASTL1 | 3625 | SINE<br>AluSg | AS | LTR<br>MER41A,<br>ERV1 family | S | 60 |
| ENST00000433673.1 | AC068282.3-001 | 2690 | SINE<br>AluSx | AS |  | AS | 58 |
| ENST00000590745.1 | CYP4F8-006 | 3117 | SINE<br>AluSx | AS |  | S | 84 |

Table S3 is available as a separate file.

